# Integrated Antibody and DIA–Based Plasma Proteomics Uncover Temporal Host Responses to SARS–CoV–2 Infection

**DOI:** 10.1101/2025.11.06.687090

**Authors:** Tetsuya Fukuda, Aya Nakayama, Yoko Chikaoka, Yasuhiko Bando, Takeshi Kawamura

## Abstract

This is a single–case longitudinal study in which we monitored antibody responses and plasma proteome dynamics over a 310–day period in a single individual who underwent COVID–19 vaccination and SARS–CoV–2 infection. To evaluate COVID–19 antibody responses, human plasma samples collected over 310 days (August 18, 2021, to June 22, 2022) were analyzed by DIA–LC–MS/MS proteomics. This led to the acquisition of a quantitative protein profile and allowed validation of the temporal behavior of expressed proteins within the samples. A total of 1,502 proteins were identified from the plasma samples. Before vaccination, after the first dose, after the second dose, and during the period after contracting the novel coronavirus, protein quantification values during each event interval were compared. Despite minimal changes before and after COVID–19 vaccination, notable proteins exhibiting distinct high–expression and low–expression patterns were identified after SARS–CoV–2 infection.

## 2. Background

The novel coronavirus (COVID–19), discovered in 2019, has demonstrated formidable infectiousness and rapidly evolved into a global pandemic, spreading across the world and causing many infections and fatalities. Researchers worldwide have responded swiftly, focusing on the rapid development of treatments and vaccines. The approved medications were also quickly put into practical use and introduced onto the market. However, like other viruses, COVID–19 has continued to generate SARS–CoV–2 variants. Although these variants tend to exhibit reduced virulence, the overall situation remains uncertain, underscoring the need for continued vigilance [1–4].

Since the early stages of the COVID–19 pandemic, our research group has been monitoring endogenous antibody responses to SARS–CoV–2 in human subjects and has collected and stored a large number of clinical plasma samples. In–depth proteomic analysis can be performed on the obtained plasma samples, particularly those taken before and after COVID–19 vaccination and infection. This can reveal how infection progresses through the variations and behaviors of specific proteins indicative of immune responses, particularly those proteins with fluctuating profiles.

In this study, blood samples were collected from a single research volunteer prior to the administration of the COVID–19 vaccination. Sequential sampling was conducted, covering the first vaccine dose (Moderna), the second dose (Moderna), subsequent COVID–19 infection, and ongoing periodic sampling thereafter. Antibody testing was carried out on the obtained samples, along with the measurement and evaluation of antibody titers. Additionally, a time series protein profile was obtained through proteomic analysis.

For proteomic analysis via LC–MS, the DIA–LC–MS method was applied [5–8]. DIA– LC–MS differs from conventional LC–MS proteomic analysis, in which only peptide– like spectra are selected from the MS spectrum of single MS in the first stage and then subjected to successive MS/MS. LC–MS proteomic analysis thus has limitations in terms of the number of MS/MS events due to the performance of the mass spectrometer. Weak peaks and similar components may be overlooked as a result. In contrast, with DIA–LC–MS, in DIA mode, for example, by segmenting the MS range, such as m/z 425–445, into segments of m/z 20 each, all the observed peaks in that range are collectively subjected to MS/MS. As a result, the identification count tends to increase compared with that of conventional shotgun LC–MS proteomic analysis.

In addition to antibody testing, we conducted DIA–LC–MS–based proteome analysis on the same plasma samples (Figure 1). Through this process, a quantitative protein profile was obtained, and the behavior of proteins expressed within the sampled time series was verified. In this report, we provide an account of our findings.

**Figure 1.**
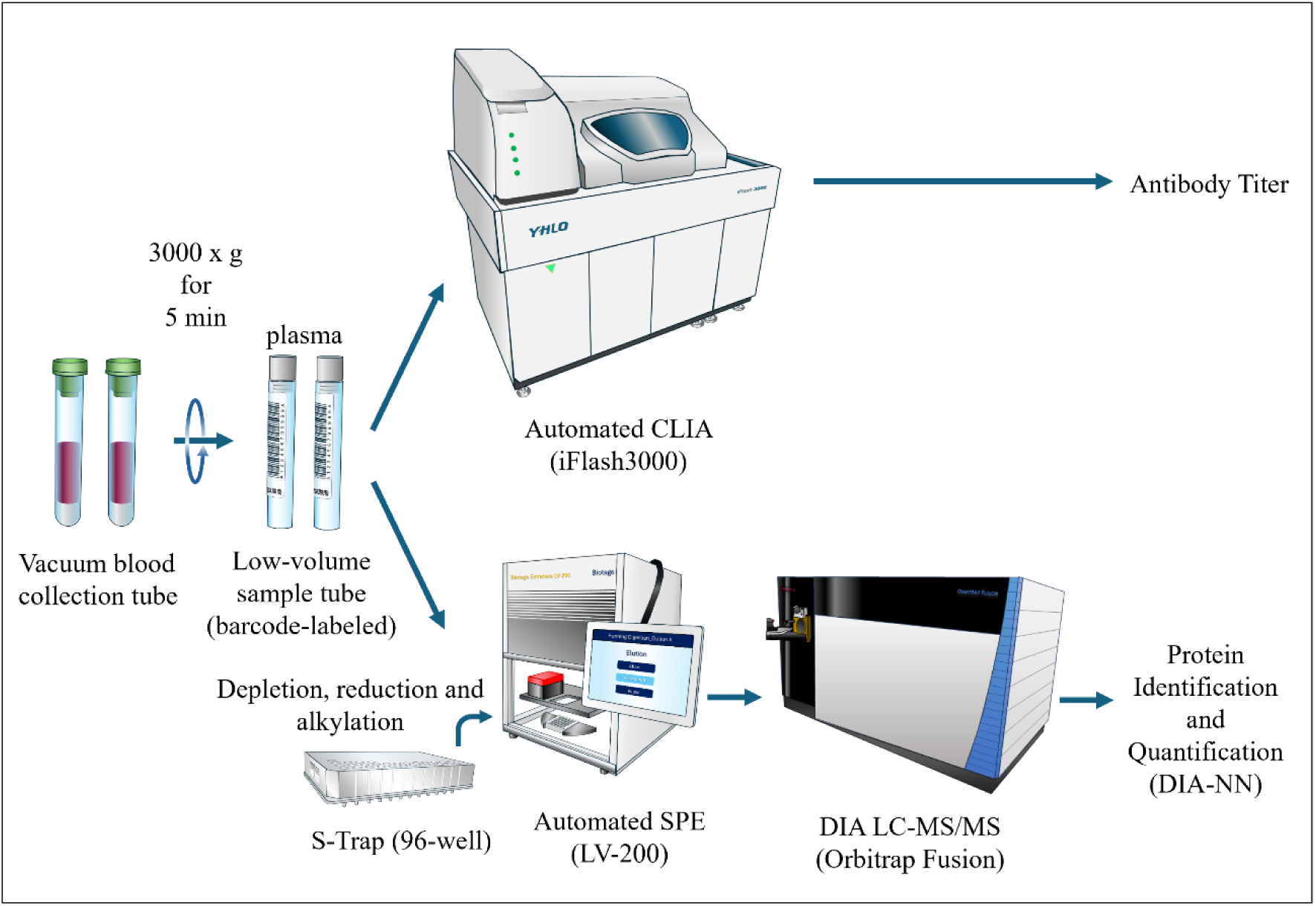
Overview of the experimental workflow for plasma proteomics and antibody analysis.

## 3. Methods

### 3.1. Samples

An overview of the entire experimental workflow is shown in Figure 1. The samples used in this study consisted of 18 human plasma samples collected from a single volunteer donor through blood sampling between August 18, 2021, and June 22, 2022. A 54–year–old male volunteer with well–controlled mild bronchial asthma participated in this study. He was on regular medication for asthma management but had no other chronic illnesses and maintained overall good health. He had received two doses of mRNA–based COVID–19 vaccination, with the last dose administered six months before infection onset. The management of these samples was conducted in accordance with institutional guidelines (The University of Tokyo) to ensure anonymity. The research involving these samples was approved by the Ethics Committee established at the University of Tokyo (Approval Number: 23–230).

Peripheral blood samples were collected in vacuum blood collection tubes. After centrifugation, plasma was separated and aliquoted into barcode–labeled low–volume tubes (φ13×75 mm). Two parallel analyses were performed from the same plasma sample:

1. Antibody profiling: Plasma was subjected to automated chemiluminescent immunoassay (CLIA) using the iFlash 3000 (YHLO) to measure IgG antibody titers against SARS–CoV–2 antigens.
2. Proteome analysis: Proteins were extracted using the Extrahera LV–200 automated solid–phase extraction system (Biotage, Uppsala, Sweden), followed by sample preparation using the S–Trap (96–well) protocol. In this study, stringent SDS removal was performed during the S–Trap cleanup process using the Extrahera LV–200: SDS– containing samples were trapped in the S–Trap column and washed five times with 90% methanol/100 mM TEAB buffer (pH 7.1) to ensure complete SDS elimination. The resulting peptides were analyzed by data–independent acquisition (DIA) using Orbitrap Fusion LC–MS/MS (Thermo Fisher Scientific), and protein identification and quantification were performed using DIA–NN software.

### 3.2. Reagents

Protease inhibitor (Protease Inhibitor Cocktail Tablets, complete, Mini, EDTA–free Tablets) was purchased from Roche Diagnostics (Indianapolis, USA). Benzonase (Benzonase® endonuclease, purity grade I (≥ 99%) suitable for biopharmaceutical production) was obtained from Merck (Darmstadt, Germany). Phosphate–buffered saline (PBS), ammonium bicarbonate (AMBIC), dithiothreitol (DTT), iodoacetamide (IAA), and triethylammonium bicarbonate (TEAB) were acquired from Sigma (St. Louis, MO, USA). For the depletion of plasma samples, the Albumin and IgG Removal Kit (Amersham Biosciences, UK) was used. Sequencing–grade trypsin was purchased from Promega (Madison, WI, USA). BCA reagent was obtained from Thermo Fisher Scientific (Pierce Biotechnology, Madison, WI, USA). Acetonitrile, trifluoroacetic acid, formic acid, and methanol were procured from Kanto Chemical Co. Ltd. (Tokyo, Japan). SDS was purchased from Amersham Biosciences (Amersham, UK). Phosphoric acid was obtained from Nacalai Tesque, Inc. (Kyoto, Japan). All water used in this study was produced via a Milli–Q system (Merck Millipore, Billerica, MA, USA).

For S–Trap processing, the following buffers and reaction solutions were prepared: 5% SDS in 50 mM TEAB (pH 7.55), 12% phosphoric acid, 90% methanol in 100 mM TEAB (pH 7.1), 50 mM TEAB (pH 8), 0.2% formic acid, and 0.2% formic acid in 50% ACN.

### 3.3. Antibody assay using iFlash 3000

In parallel with the proteomic analysis, plasma antibody titers against SARS–CoV–2 were measured. Blood samples were collected into vacuum blood collection tubes (for plasma separation), followed by centrifugation at 3,000 rpm for 5 minutes. The supernatant plasma was carefully recovered and subjected to a chemiluminescence immunoassay (CLIA).

Plasma antibody titers against SARS–CoV–2 were quantified using the fully automated chemiluminescent immunoassay (CLIA) analyzer iFlash 3000 (Shenzhen YHLO Biotech Co., Ltd., China). The assays included IgG antibodies targeting the nucleocapsid (N), spike subunit 1 (S1), and receptor–binding domain (RBD) antigens, as well as neutralizing antibodies (NAbs). All measurements were carried out with manufacturer– supplied reagent kits, applying a positivity threshold of 10 AU/mL.

The iFlash system employs paramagnetic microparticle–based CLIA technology, enabling high–throughput and multiplexed detection. The neutralization assay is based on a surrogate virus neutralization test (sVNT), which evaluates the inhibition of ACE2 binding to RBD–coated particles and thereby provides functional information on antibody–mediated neutralization. Calibration was performed using lot–specific QR code–encoded master curves with dedicated calibrators (CAL1 and CAL2 or CAL1– CAL4, depending on the assay), and internal quality controls at two concentration levels were included in each run according to the manufacturer’s instructions (IFU, version 5.0, 2020). The reproducibility of the iFlash assays has been validated to ≤8% CV, with high sensitivity and specificity demonstrated in previous evaluations (Figure 1) [9–13].

### 3.4. Sample preparation for DIA–LC–MS

For proteomic analysis via DIA–based LC–MS, plasma samples were processed as follows (Figure 1).

The major proteins in plasma, such as albumin and IgG, were removed using the Albumin and IgG Removal Kit (Amersham Biosciences, UK) as follows:

First, the resin in the vial was thoroughly mixed to create a suspension, and an appropriate amount (approximately 5 mL) was transferred to a 15 mL tube. The resin was subsequently washed three times with PBS. A total of 15 μL of the plasma sample was added to 750 μL of the resin slurry, and the mixture was shaken at room temperature for 30 min (1400 rpm, rt–21°C, 30 min → manual shaking at 250 rpm). The mixture was then subjected to centrifugation using a spin column (6500 × g, 5 min), and the flowthrough was collected. The BCA protein assay was conducted to quantify the protein amount, and the results revealed a total protein quantity of 2.1–2.3 mg.

Next, TCA–acetone precipitation was performed, and the collected precipitate was dissolved in 5% SDS in 50 mM TEAB (pH 7.55), resulting in a total volume of 50 μL. Subsequently, reduction and alkylation of the sample were carried out. Then, 1 M DTT was added to achieve a final concentration of 20 mM, followed by incubation at 95°C for 10 min and then a return to room temperature. Next, 1 M IAA was added to achieve a final concentration of 40 mM, and the sample was incubated in the dark at room temperature for 30 min. After reduction and alkylation, 12% phosphoric acid was added to the sample mixture to achieve a final concentration of 1.2%. A total of 350 μL of 90% methanol in 100 mM TEAB (pH 7.1) was added to bring the total volume to 405 μL. The sample was then subjected to S–Trap 96–well processing. The sample was gently transferred to S–Trap 96–well plate, to which 250 μL of 90% methanol in 100 mM TEAB (pH 7.1) was added for protein retention and subsequent washing of the retained proteins. This operation was repeated four times consecutively. Extrahera LV–200 (Biotage, Uppsala, Sweden) was used for these steps (Figure 1).

Then, 250 μL of 50 mM TEAB was added to 20 μg/vial of trypsin (Promega) for dissolution, and 125 μL of this solution was immediately added to the S–Trap 96–well, which was then centrifuged at 4000 × g for 1 min. After incubation at 47°C for 60 min, the reaction was continued overnight at 37°C. The S–Trap 96–well plate was set on the Extrahera LV–200, and 80 μL of 50 mM TEAB was added to recover the flowthrough. Subsequently, 80 μL of 0.2% HCOOH was added, and the flowthrough was collected. Furthermore, 80 μL of 0.2% HCOOH in 50% ACN was added, followed by centrifugation at 4000 × g for 1 min to separate the flowthrough, which was then collected. After drying using a SpeedVac, the recovered samples were reconstituted in 200 μL of 0.1% TFA in 2% ACN.

### 3.5. LC–MS/MS conditions

The eluted samples were separated by nanoflow reverse–phase LC followed by analysis via an Orbitrap Fusion mass spectrometer (Thermo Fisher Scientific, San Jose, CA, USA) equipped with a Dream spray nano–electrospray ionization source (AMR Inc., Tokyo, Japan) [14–19]. The LC used was an Advance UHPLC (ultrahigh–performance liquid chromatography) instrument (Bruker Daltonics, Bremen, Germany) equipped with an HTS PAL auto–sampler (CTC Analytics AG, Zwingen, Switzerland). The samples were loaded onto a capillary reverse–phase separation column packed with 3.0–μm–diameter gel particles with the pore size 120 Å (L–column2 ODS, 3 μm, 0.2 × 150 mm; CELI, Tokyo, Japan). Eluent A was 0.1% formic acid, and eluent B was 100% acetonitrile. The column was eluted at a flow rate of 1.2 μL/min with a concentration gradient of A + 5% B to 35% B in 100 min and 35% B to 95% B in 1 min. This was followed by isocratic elution with 95% B for 8 min and a further concentration gradient from 95% B to 5% B in 1 min [20].

The mass spectrometer was operated in data–independent acquisition (DIA) mode, in which MS acquisition with a mass range of m/z 495–745 was automatically switched to MS/MS acquisition under the automated control of Xcalibur software 3.1 (Thermo Fisher Scientific).

DIA data were collected via staggered windows with a loop count of 20.0 m/z isolation windows, and the collision energy mode was stepped with 5 ms maximum injection time in custom mode.

In DIA mode, each cycle consisted of a MS1 scan of 495–745 m/z with resolution of 60,000 and AGC target of 2 × 10^6^, followed by 50 MS2 scans of 498–742 m/z with resolution of 30,000 and AGC target of 2.5 × 10^6^ [7,8,21].

### 3.6. DIA data processing

Raw data were processed using Data–Independent Acquisition by Neural Networks (DIA–NN) for proteome analysis. DIA–NN is known to be particularly useful for applications in high–throughput proteomics because it improves the performance of protein identification and quantification in conventional DIA–mode proteomics applications, enabling fast and reliable protein identification [6,22].

The algorithm for DIA–NN was as follows. DIA–NN version 1.8.1, in library–free mode, was used with the same UniProt FASTA database. The protein database was the UniProt Reference Proteome for Homo sapiens (Taxonomy 9606; Proteome ID UP000005640; 20,373 entries; UniProt release 2022_03; reviewed human canonical).

Precursors of charge states 1–4, peptide lengths 7–30, and peptide m/z 300–1800 were considered, with a maximum of one missed cleavage. A maximum of one variable modification per peptide was considered. Cysteine carbamidomethylation was allowed as a fixed modification, N–terminal methionine excision as a variable modification, methionine oxidation as a variable modification, and N–terminal acetylation as a variable modification. The precursor false discovery rate was then filtered at 1% FDR [6–8, 21–22].

The raw and processed mass spectrometry data have been deposited in JPOST (JPST003696) and registered with ProteomeXchange (PXD062242).

### 3.7. Multivariate and differential expression analysis

To evaluate temporal protein expression dynamics, multivariate and differential expression analyses were conducted using Scaffold DIA (version 3.4.1, Proteome Software, Portland, OR, USA). DIA–MS sample data were loaded into Scaffold DIA from search engine result files. Protein identifications were grouped according to the principle of parsimony; those that could not be distinguished by MS/MS spectra were clustered, and filtered to achieve a protein–level false discovery rate (FDR) of less than 1.0%.

Differentially expressed proteins were identified based on fold–change calculations, and proteins showing a fold change ≥ 1.9 after SARS–CoV–2 infection were selected as the Top 32 proteins. Principal Component Analysis (PCA) was performed to visualize global variance across samples, and hierarchical clustering heatmaps were generated using normalized protein intensity values. Volcano plots were used to identify significantly upregulated or downregulated proteins between the pre– and post–infection groups (A and B), incorporating both fold change and p–value thresholds.

All calculations and visualizations (PCA, Heatmap, Volcano plot) were conducted within the Scaffold DIA environment. Gene ontology (GO) annotations were not applied in this analysis. The statistical significance of peptide identifications was assessed using the Percolator algorithm and decoy database strategy. The software also utilized modules from ProteoWizard and other cited components (Chambers et al., 2012; Käll et al., 2007, 2008).

## 4. Results

### 4.1. Antibody Titer Dynamics Following Vaccination and Infection

The results of antibody titers measured using the iFlash 3000 are summarized in Table 1 and Figure 2. As shown in the graph, both the S1 antigen and neutralizing antibody values increased markedly after the second vaccination. In contrast, the IgG antibody against the N antigen did not increase in response to vaccination. However, a sharp rise in the IgG N antigen antibody value was clearly observed following SARS–CoV–2 infection. Based on these findings, we proceeded to examine the proteomic profiles of the plasma samples using DIA–based analysis to identify proteins that showed a clear response associated with these antibody dynamics.

**Table 1.**
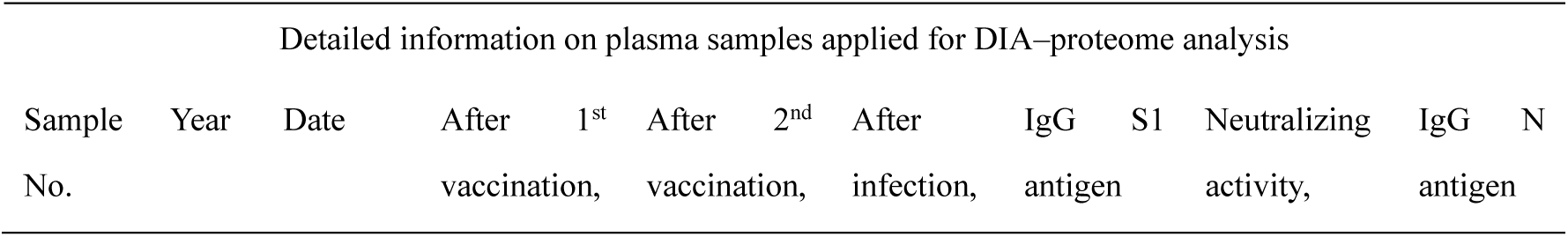
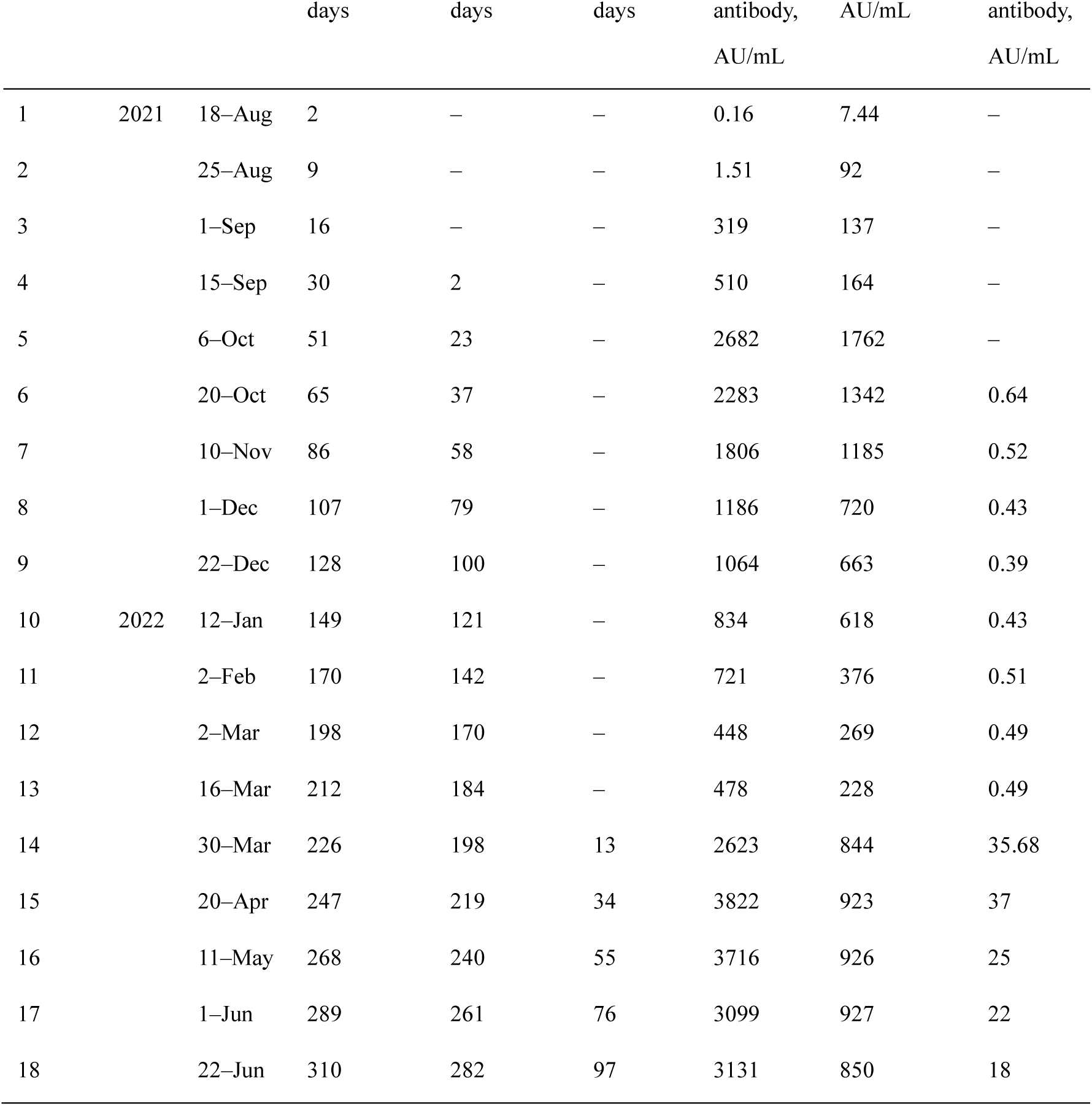
Summary of antibody titers, collection dates, and DIA–based proteome analysis of human plasma samples. 1st Vaccination: August 16, 2021 (Moderna), 2nd Vaccination: September 13, 2021 (Moderna), COVID–19 infection: March 17, 2022 (PCR positive), March 23, 2022 (PCR negative).

**Figure 2.**
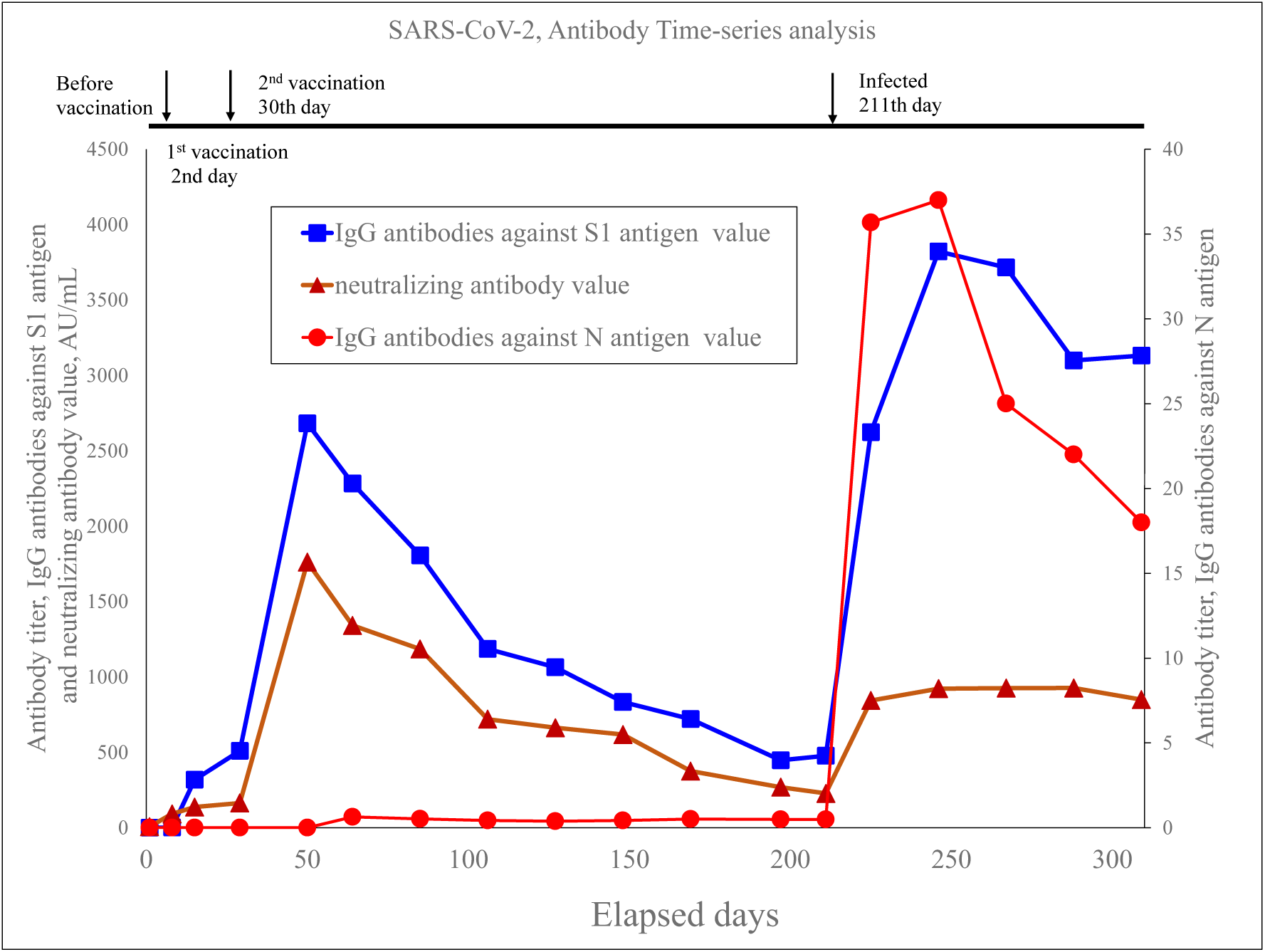
Fluctuations in the antibody titers against SARS–CoV–2 in human plasma samples.

Antibody titers in human plasma samples from a single donor were measured over a period of 310 days, from August 18, 2021, to June 22, 2022. The donor received their first dose of the vaccine (Moderna) two days after the initial blood collection, followed by the second dose 30 days later (Moderna). At 211 days after the sample collection date, the donor became infected with COVID–19.

Building on the distinct antibody titer dynamics observed before and after vaccination and infection, we conducted a comprehensive plasma proteome analysis using data– independent acquisition (DIA)–based LC–MS/MS. This approach aimed to capture both qualitative and quantitative changes in plasma proteins corresponding to individual immune responses. Following protein identification and quantification, multivariate statistical analyses including principal component analysis (PCA) and heatmap clustering were performed to visualize longitudinal trends and identify characteristic protein expression patterns linked to these immunological events. These data provided a broader molecular context that complemented the antibody titer profiles and enabled the identification of potential biomarkers reflecting the host’s systemic response to SARS– CoV–2 exposure.

### 4.2. Comprehensive multivariate analysis of plasma proteomes before and after SARS– CoV–2 infection

A total of 1,502 human proteins were identified from 18 longitudinal plasma samples using DIA–based LC–MS/MS. To understand the overall protein expression dynamics in relation to vaccination and infection events, multivariate statistical analyses were performed on the quantified proteomic data.

Principal component analysis (PCA) revealed a clear temporal shift in the global proteome profile, particularly distinguishing post–infection samples from pre– vaccination and post–vaccination ones (Figure 3). The score plot showed that samples clustered along a trajectory corresponding to the volunteer’s clinical course, with samples collected after infection forming a distinct group. This trend was further supported by hierarchical clustering of the top 200 most variable proteins, which grouped post– infection samples separately from earlier time points (Figure 4).

**Figure 3.**
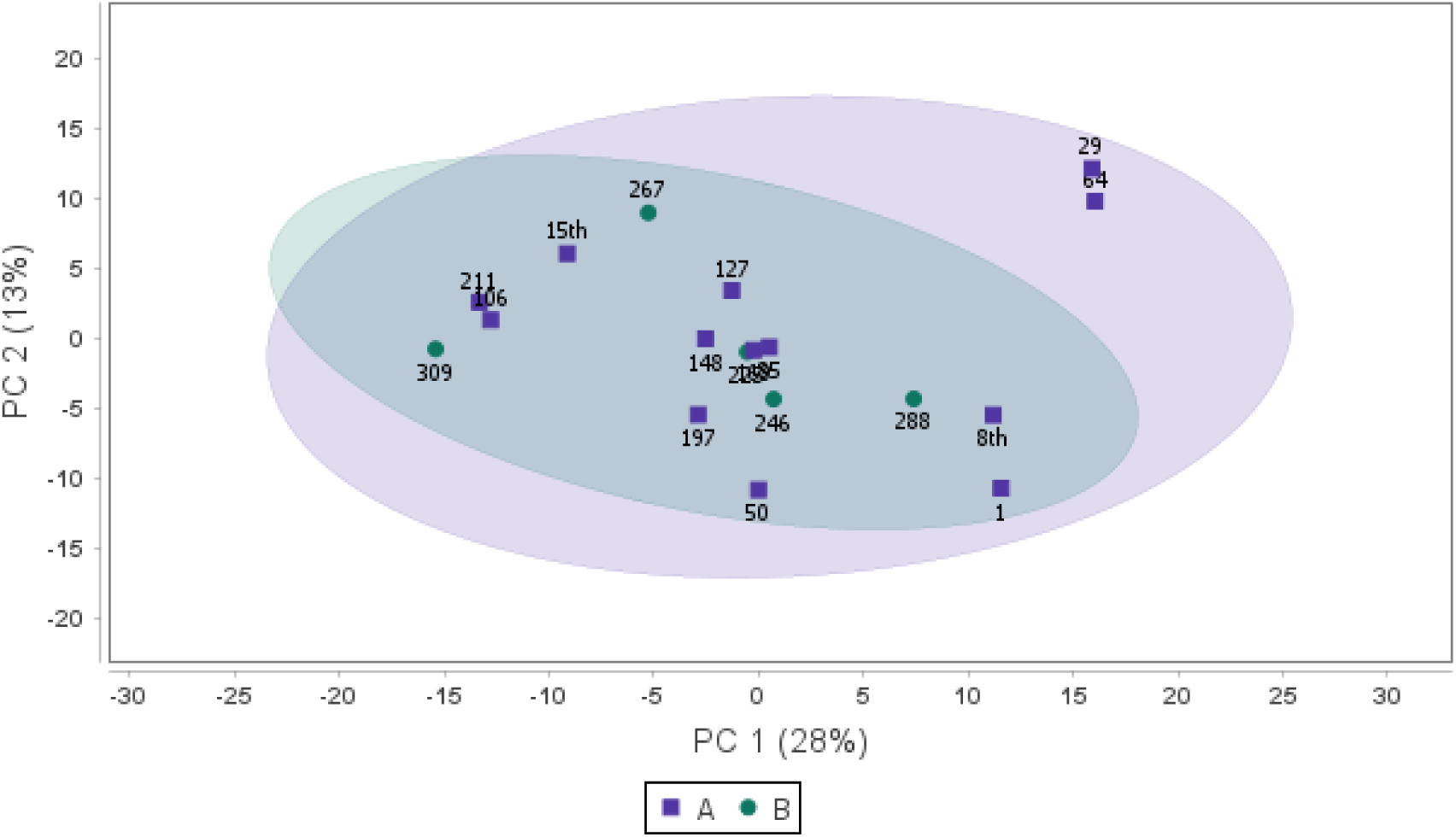
Scores plot of Principal Component Analysis (PCA) using quantified protein expression data from plasma samples.

**Figure 4.**
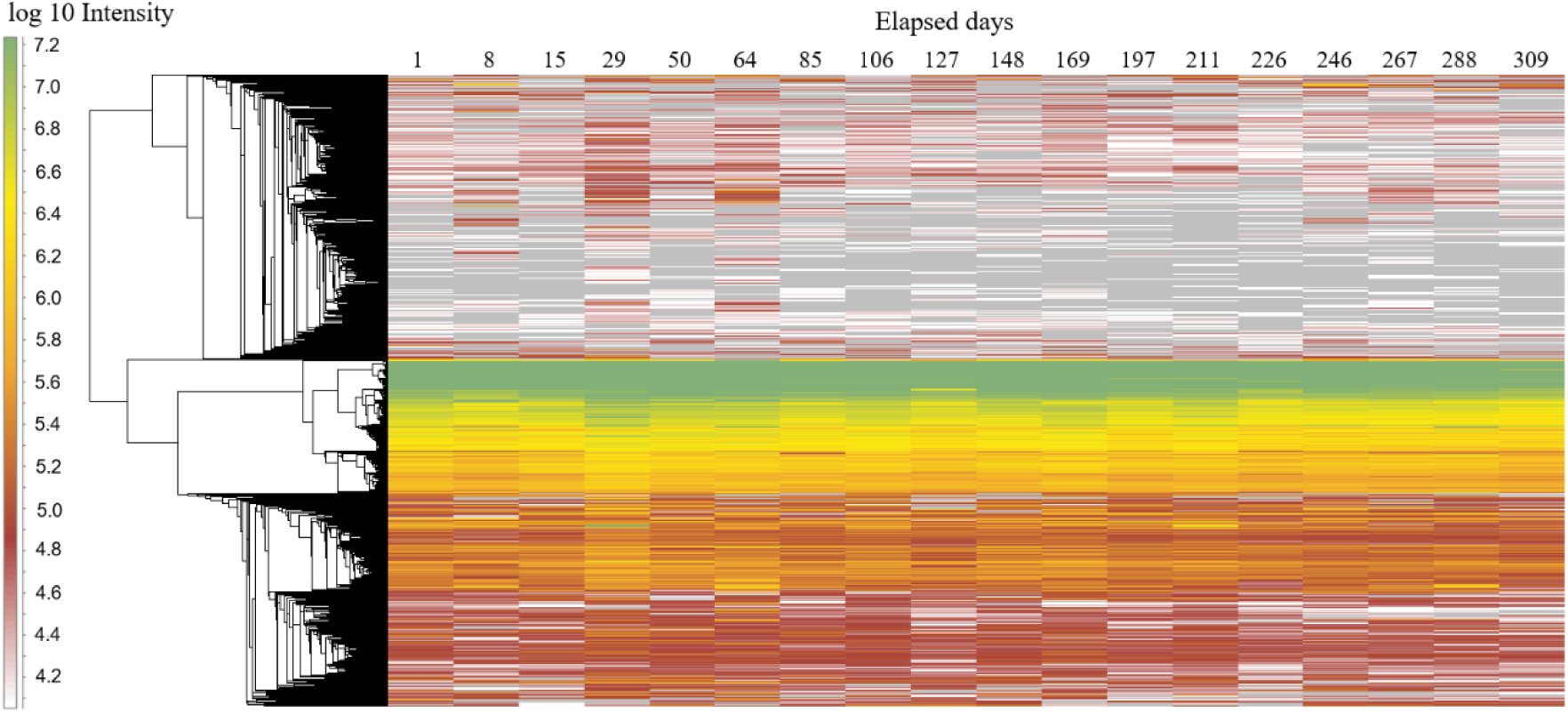
Heatmap depicting the quantitative expression profiles of all proteins identified in longitudinal plasma samples using DIA–based proteome analysis.

Each point represents an individual sample, with purple squares (▪) indicating pre– infection samples (Group A) and green circles (●) representing post–infection samples (Group B). The first two principal components, PC1 (28%) and PC2 (13%), explain 41% of the total variance. While the two groups partially overlap, a moderate shift in the distribution of Group B toward the negative side of PC1 suggests systematic proteomic changes associated with SARS–CoV–2 infection.

These findings suggest that SARS–CoV–2 infection induces marked proteomic remodeling in the plasma, beyond the changes observed following vaccination. Based on these multivariate analyses, we next aimed to identify individual proteins that showed the most prominent upregulation after infection, using fold–change–based filtering.

### 4.3. Identification of the Top 32 Most Upregulated Plasma Proteins Following SARS– CoV–2 Infection

To further visualize the longitudinal changes in plasma proteins, a heatmap analysis was performed for all proteins identified by DIA–NN. The heatmap clearly shows a marked shift in expression patterns after SARS–CoV–2 infection (sample No.14 onward), while profiles before infection and after vaccination remained relatively stable (Figure 4).

Each column represents a time point, and each row corresponds to a protein. Values are shown as log–transformed intensities. Rows are clustered using hierarchical clustering (dendrogram shown on the left). Color scale represents log10–transformed protein intensities. The overall proteomic profile appeared largely stable over the 310–day study period. However, a subset of proteins showed noticeable expression changes, particularly after SARS–CoV–2 infection (from sample No.14 onward), suggesting potential biological responses despite the overall stability.

To further identify key molecular events associated with SARS–CoV–2 infection, we extracted the top 32 plasma proteins that exhibited the highest upregulation based on fold change (FC ≥ 1.9) in samples collected after infection. This subset was selected by comparing proteomic profiles before infection (samples 1–13) with those after infection (samples 14–18), as determined by DIA–based quantification.

The selected proteins are listed in Table 2, and their expression dynamics are visualized in Figure 5 as a line graph. This graph highlights temporal increases in expression corresponding to the infection period. Several immune–related and inflammation– associated proteins were included in the top–ranked group, suggesting a strong host response to viral invasion. Notably, proteins such as CNOT1, RPL11, and GSDMA showed consistent upregulation after infection and aligned well with changes in N antigen antibody titers.

**Table 2.**
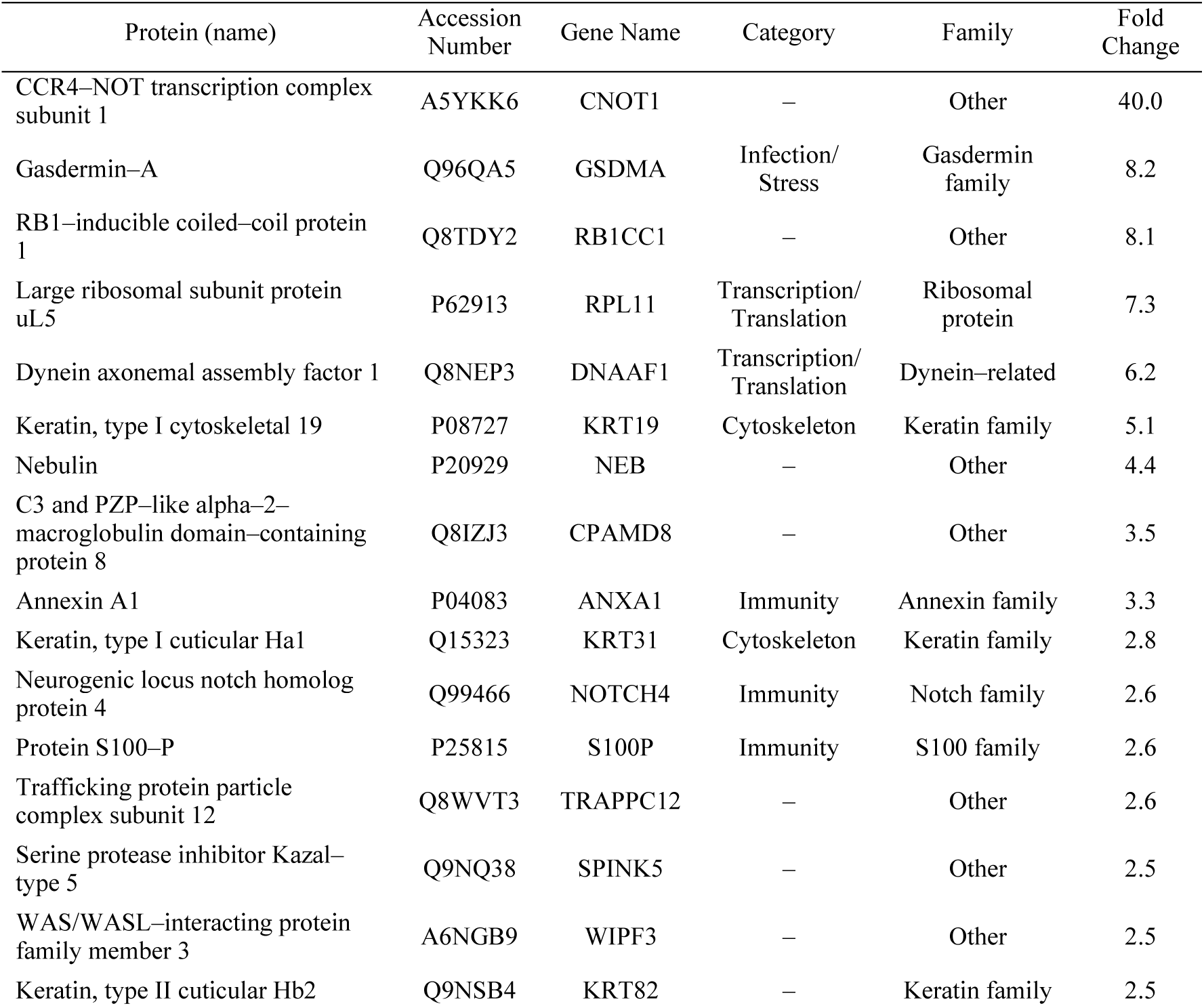
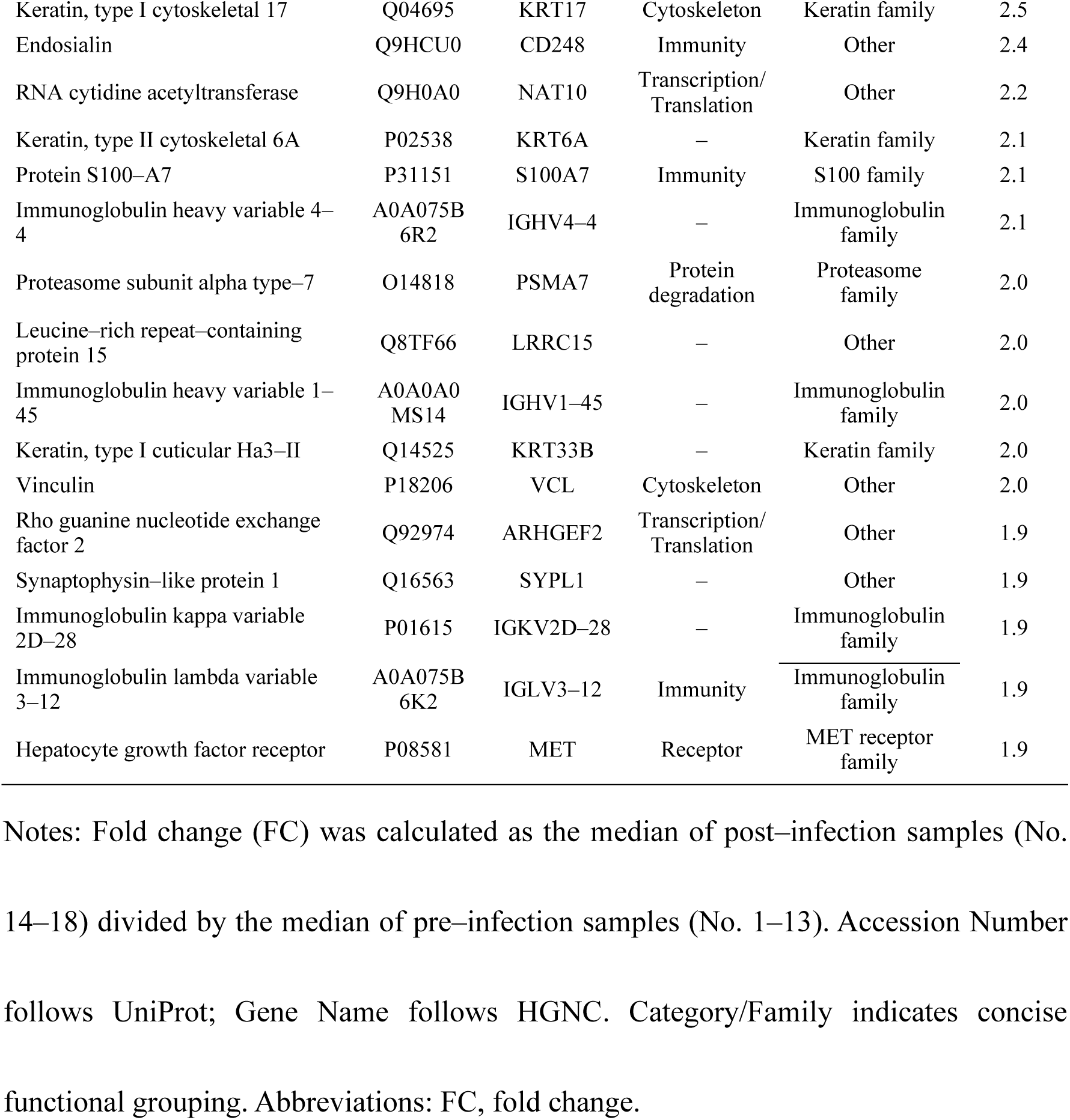
Top 32 Plasma Proteins Showing Increased Abundance Following COVID–19 Infection. This table lists the top 32 plasma proteins that showed increased abundance after COVID–19 infection, as identified by DIA–NN analysis. Proteins were selected based on a fold change (infection vs. pre–infection) ≥ 1.9. For each protein, the function or interpretation, UniProt accession number, gene name, functional category, protein family, and fold change are provided. Categories include immunity–related proteins, cytoskeletal components, and transcription/translation–associated proteins, among others. This classification aids in understanding the biological relevance of the proteomic changes observed in the longitudinal study. Notes: Fold change (FC) was calculated as the median of post–infection samples (No. 14–18) divided by the median of pre–infection samples (No. 1–13). Accession Number follows UniProt; Gene Name follows HGNC. Category/Family indicates concise functional grouping. Abbreviations: FC, fold change.

**Figure 5.**
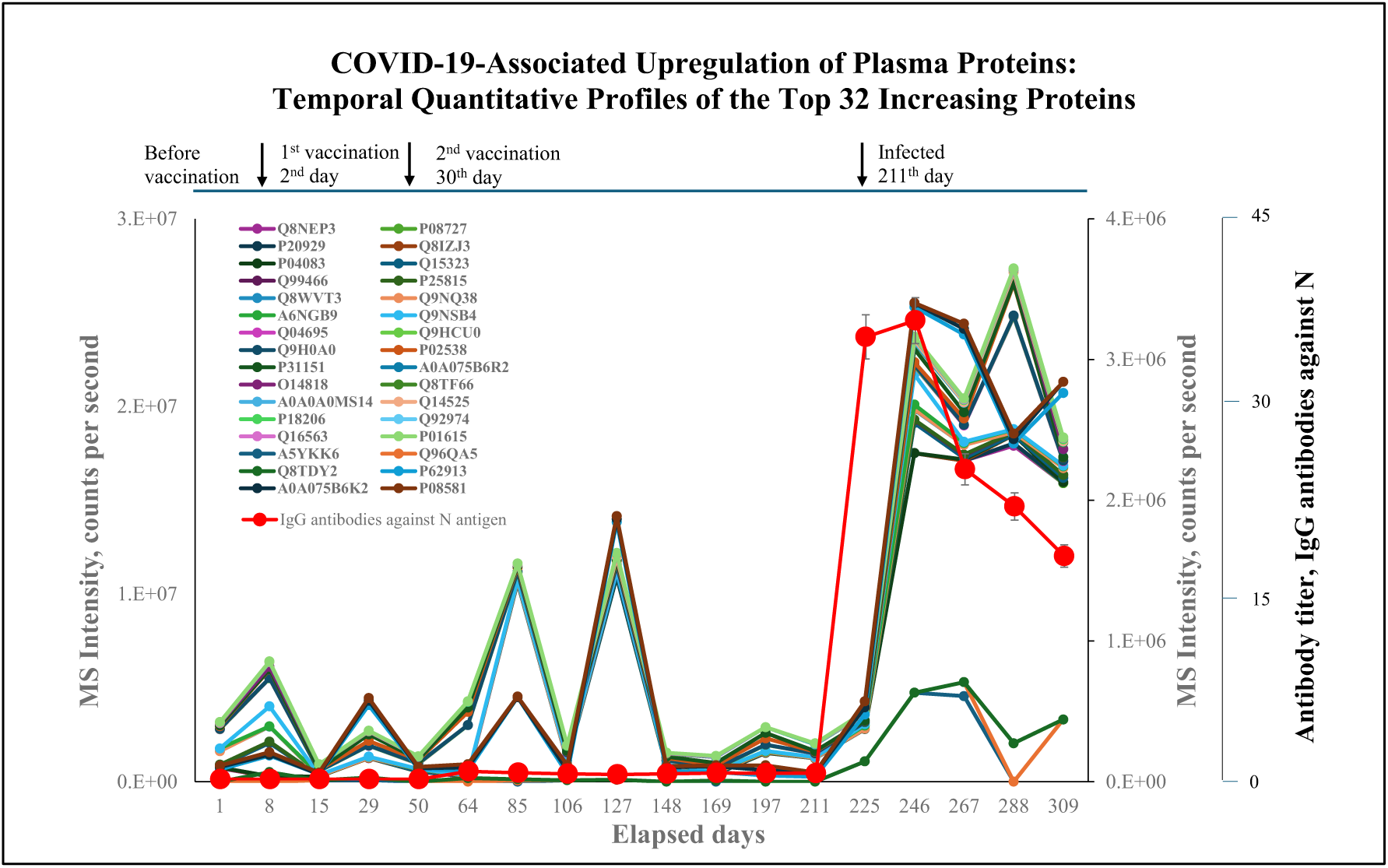
Temporal expression profiles of the top 32 upregulated plasma proteins following SARS–CoV–2 infection.

Proteins were selected from all quantified proteins based on a fold change ≥ 1.9, as determined by DIA–NN analysis. The line graph shows the MS intensity (counts per second) of each protein across 18 time points over a 310–day period. The red line indicates IgG antibody titers against the SARS–CoV–2 N antigen, measured using the iFlash 3000 (right axis). A sharp increase in both IgG levels and MS intensities of selected proteins was observed after COVID–19 infection on day 211, whereas minimal changes were seen before and after vaccination.

These results reinforce the potential utility of DIA–MS in capturing biologically relevant proteomic shifts during immune responses and provide candidate markers for future investigation into infection–induced pathways.

### 4.4. Volcano Plot–Based Screening of Significantly Regulated Proteins

To statistically identify proteins that were significantly regulated after SARS–CoV–2 infection, we generated a volcano plot based on the DIA–MS quantitative data (Figure 6). The analysis compared samples collected before infection (samples 1–13) and after infection (samples 14–18), applying a fold change (FC) threshold of >1.5 and a p–value cutoff of <0.05.

**Figure 6.**
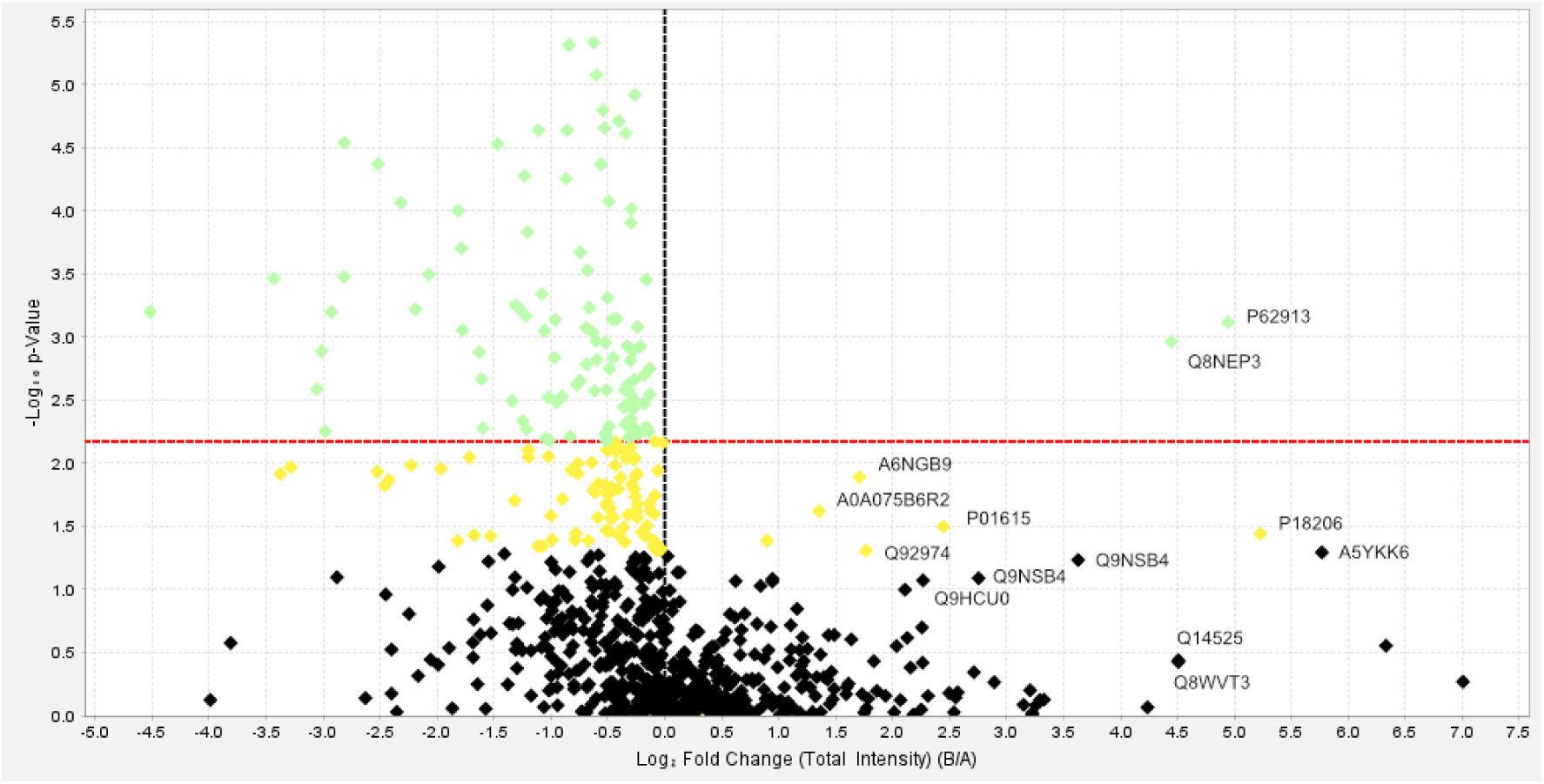
Volcano plot showing differentially expressed plasma proteins before and after SARS–CoV–2 infection.

Group A represents pre–infection samples, and Group B represents post–infection samples. The horizontal axis shows log₂ fold changes (B/A), and the vertical axis shows –log₁₀(p–values). Proteins with increased expression after infection are plotted on the right side. Among the top 32 most upregulated proteins (fold change ≥ 1.9), 12 representative proteins (A5YKK6, P62913, Q8NEP3, Q8WVT3, A6NGB9, Q9NSB4, Q9HCU0, A0A075B6R2, Q14525, P18206, Q92974, P01615) are labeled in the plot.

These were selected based on their distinct positioning and clarity in the plot, as the remaining proteins were in a densely clustered region where labeling was visually impractical.

As shown in Table 2 and Figure 5, multiple proteins were identified as significantly upregulated in the post–infection group. Proteins such as CNOT1, RPL11, and GSDMA showed both high statistical significance and large fold changes, confirming their role as potential markers of infection–associated immune responses. Conversely, proteins such as U1SBP (Q16560) were downregulated and may reflect suppressed pathways or altered transcriptional activity following infection (Supplementary Figure 1).

The volcano plot visualization provided a clear and intuitive representation of the global proteomic changes induced by SARS–CoV–2 infection and allowed for the prioritization of proteins for downstream validation and mechanistic exploration.

### 4.5. Proteomic Characterization Based on the Top 32 Most Upregulated Proteins

To explore proteomic trends associated with the host response to SARS–CoV–2 infection, we extracted the top 32 most upregulated plasma proteins based on fold change values comparing post–infection samples (No. 14–18) to pre–infection samples (No. 1–13). The selected proteins exhibited fold changes ≥ 1.9 and encompassed a broad range of functional categories (Table 2).

Notably, these proteins included molecules associated with transcription/translation (e.g., CNOT1, RPL11), cytoskeletal regulation (e.g., KRT19, KRT17, NEB), immune responses (e.g., ANXA1, S100P, NOTCH4, IGHV4–4), and protein degradation (e.g., PSMA7). The diversity of protein classes suggests a systemic biological response, including inflammation, immune activation, structural remodeling, and potential epigenetic regulation.

Among these, the 60S ribosomal protein L11 (RPL11) and CCR4–NOT complex subunit 1 (CNOT1) showed particularly high fold changes (>7.0 and 40.0, respectively), indicating marked upregulation of the translational machinery. Also of note were S100P and Gasdermin–A, implicating inflammatory and pyroptotic responses.

These observations suggest that SARS–CoV–2 infection induces broad molecular alterations beyond conventional immunoglobulin responses, potentially involving post– transcriptional and cytoskeletal remodeling processes. Further mechanistic validation is warranted, and detailed interpretation of these changes is discussed in the following section (Discussion).

## 5. Discussion

In this study, we demonstrated that the parallel application of SARS–CoV–2 antibody testing and DIA–based LC–MS/MS proteome profiling to the same plasma samples enabled time–resolved visualization of protein dynamics associated with antibody titer fluctuations. This integrative approach led to the identification of candidate biomarkers and provided insight into host responses to SARS–CoV–2 infection.

Comprehensive multivariate analysis–including PCA, heatmaps, and volcano plots– highlighted distinct post–infection expression patterns. In particular, the upregulation of 60S ribosomal protein L11 (RPL11) and CCR4–NOT transcription complex subunit 1 (CNOT1) was striking. RPL11, a structural component of the large ribosomal subunit, is also involved in p53–mediated stress responses and ribosomal biogenesis regulation [26–31]. These findings are consistent with previous studies reporting increased expression of RPL29 and RPL18 in SARS–CoV–2 models, supporting the hypothesis that viral infection enhances translational activity via ribosomal machinery [23–25].

Notably, CNOT1 exhibited the highest fold change (>40–fold) among all proteins. As a scaffold protein of the CCR4–NOT complex, CNOT1 regulates mRNA deadenylation, transcriptional repression, and immune signaling. Its dramatic induction implies activation of post–transcriptional gene regulation following infection.

In addition to translational machinery components, we also observed elevated expression of immune modulators (e.g., ANXA1, S100P, NOTCH4), cytoskeletal proteins (e.g., KRT17, KRT19), and inflammation–related factors (e.g., GSDMA). These changes suggest a broad host response involving inflammation, immune regulation, and tissue remodeling.

Conversely, some nuclear regulatory proteins such as U1SBP (Q16560) were markedly downregulated. This protein is associated with pre–mRNA splicing and transcription regulation, and its loss may imply compromised nuclear RNA processing. Such dysregulation could impair stress recovery or exacerbate disease progression, especially in immunocompromised individuals [31–34]. While beyond the scope of the present study, this observation will be prioritized in future validation studies.

Overall, this study underscores the potential of integrated antibody–proteomics approaches for deciphering host–pathogen interactions and supports further exploration of key molecular signatures such as RPL11 and CNOT1 in COVID–19 pathophysiology.

These findings suggest that combined proteome and antibody analyses may provide novel insights into host responses against SARS–CoV–2 infection and support the development of new biomarkers and therapeutic targets. However, this study has certain limitations. Neutralizing antibody assays, such as pseudovirus–based tests, were not performed, which may restrict the interpretation of functional immune responses. Addressing this point will be an important task in future investigations.

## Acknowledgements

We would like to express our sincere appreciation to Ms. Akiko Fukuda (Laboratory for Systems Biology and Medicine, Isotope Science Center, The University of Tokyo) for her invaluable assistance in conducting the antibody titer measurements, and to Ms. Noriko Kagi (Biotage Japan Ltd., Tokyo, Japan) for her expert technical support with the LV– 200 experiments.

## 6. Declarations

### 6.1. Declaration of generative AI and AI–assisted technologies in the writing process

During the preparation of this work, the author used ChatGPT (OpenAI) to improve the clarity and readability of the manuscript. After using this tool, the authors reviewed and edited the content as needed and took full responsibility for the content of the publication.

### 6.2. Ethics approval and consent to participate

This study was approved by the Institutional Ethics Committee of The University of Tokyo (approval number: 23–230). Informed consent was obtained from all individual participants included in the study.

### 6.3. Funding

This study was supported by Medical & Biological Laboratories Co., Ltd., and Shenzhen YHLO Biotech Co., Ltd. (the distributor and manufacturer of the antibody measurement system, iFlash 3000); by grants from Peace Winds Japan, Kowa Co., and the Research Center for Advanced Science and Technology at the University of Tokyo; and by the Japan Agency for Medical Research and Development (AMED) under the Grant–in–Aid for the Development of Vaccines for the Novel Coronavirus Disease (Grant Number: 22nf0101638h0002).

### 6.4. Conflicts of interest

The authors declare that they have no conflict of interest.

## Notes

### Competing Interest Statement

The authors have declared no competing interest.

