## Supplementary Figure 1 for "Integrated Antibody and DIA–Based Plasma Proteomics Uncover Temporal Host Responses to SARS–CoV–2 Infection"

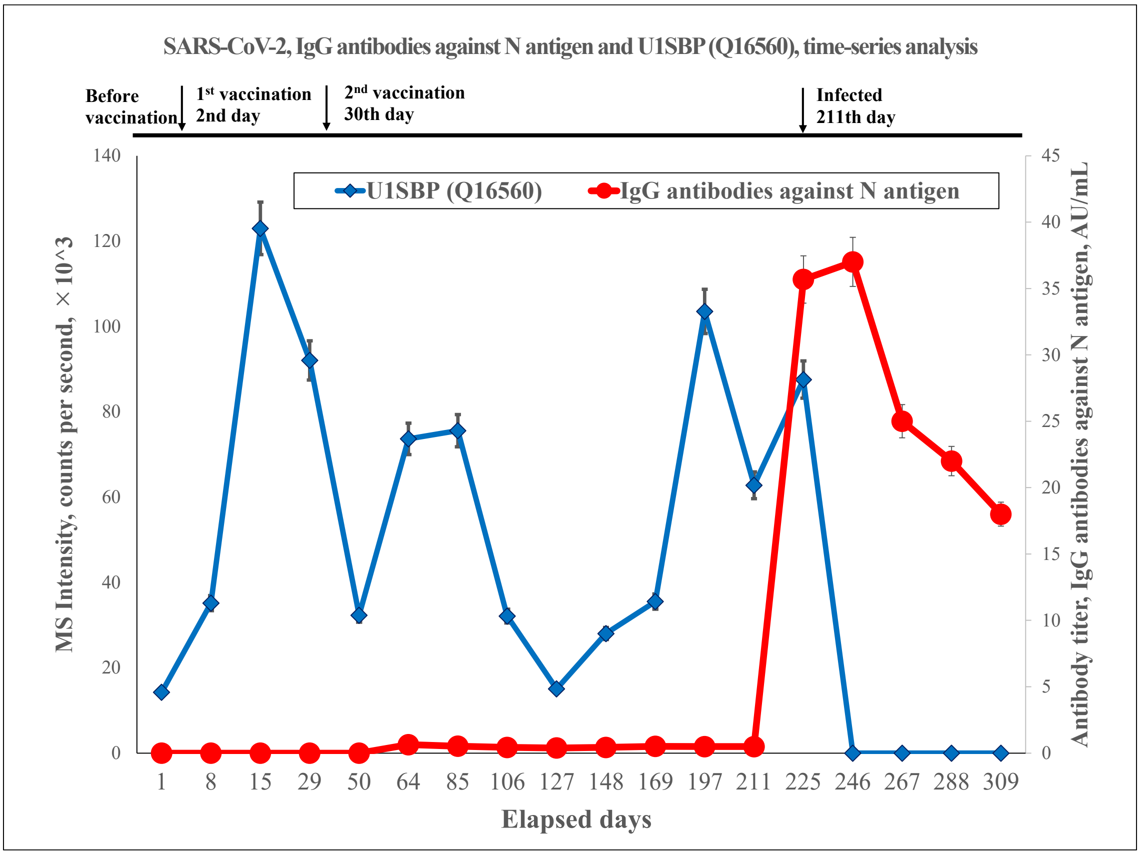


Supplementary Figure 1.

Time-series profile of U1SBP (Q16560) in plasma proteome and anti-N IgG titers.

MS intensity values for U1SBP (green line; left y-axis; counts per second, ×10^3) and IgG antibody titers against the SARS-CoV-2 N antigen (red line; right y-axis; AU/mL) were plotted across 18 time points over a 310–day period from the initial blood collection. Notably, U1SBP showed marked downregulation around SARS-CoV-2 infection (day 211), whereas anti-N IgG increased sharply at the same time point, revealing an inverse pattern. Although U1SBP was not included among the Top 32 upregulated proteins selected by fold-change analysis, this single-case observation suggests a downregulated candidate that warrants follow-up validation.
